# The relationship between transmission time and clustering methods in *Mycobacterium tuberculosis* epidemiology

**DOI:** 10.1101/302232

**Authors:** Conor J Meehan, Pieter Moris, Thomas A. Kohl, Jūlija Pečerska, Suriya Akter, Matthias Merker, Christian Utpatel, Patrick Beckert, Florian Gehre, Pauline Lempens, Tanja Stadler, Michel K. Kaswa, Denise Kühnert, Stefan Niemann, Bouke C de Jong

**Affiliations:** Unit of Mycobacteriology, Biomedical Sciences, Institute of Tropical Medicine, Antwerp, 2000, Belgium; Advanced Database Research and Modelling (ADReM), Department of Mathematics and Computer Science, University of Antwerp, Antwerp, 2020, Belgium; Biomedical Informatics Research Network Antwerp (biomina), University of Antwerp, Antwerp, 2020, Belgium; German Center for Infection Research, Partner Site Hamburg-Lübeck-Borstel-Riems, D-23845 Borstel, Germany; Molecular and Experimental Mycobacteriology, Priority Area Infections, Research Center Borstel, D-23845 Borstel, Germany; Department of Biosystems Science and Engineering, ETH Zürich, 4058 Basel, Switzerland; Vaccines and Immunity Theme, Medical Research Council Unit The Gambia, Serekunda, The Gambia; Bernhard Nocht Institute for Tropical Medicine, 20359 Hamburg, Germany; University Hospital Zurich, 8091 Zurich, Switzerland; National Tuberculosis Program, B.P 1197, Kinshasa, Democratic Republic of Congo

**Keywords:** *Mycobacterium tuberculosis*, MDR-TB molecular epidemiology, transmission, spoligotyping, MIRU-VNTR, MLST, whole genome sequencing, outbreak detection

## Abstract

**Background:** Tracking recent transmission is a vital part of controlling widespread pathogens such as *Mycobacterium tuberculosis*. Multiple methods with specific performance characteristics exist for detecting recent transmission chains, usually by clustering strains based on genotype similarities. With such a large variety of methods available, informed selection of an appropriate approach for determining transmissions within a given setting/time period is difficult.

**Methods:** This study combines whole genome sequence (WGS) data derived from 324 isolates collected 2005-2010 in Kinshasa, Democratic Republic of Congo (DRC), a high endemic setting, with phylodynamics to unveil the timing of transmission events posited by a variety of standard genotyping methods. Clustering data based on Spoligotyping, 24-loci MIRU-VNTR typing, WGS based SNP (Single Nucleotide Polymorphism) and core genome multi locus sequence typing (cgMLST) typing were evaluated.

**Findings:** Our results suggest that clusters based on Spoligotyping could encompass transmission events that occurred over 70 years prior to sampling while 24-loci-MIRU-VNTR often represented two or more decades of transmission. Instead, WGS based genotyping applying low SNP or cgMLST allele thresholds allows for determination of recent transmission events in timespans of up to 10 years e.g. for a 5 SNP/allele cut-off.

**Interpretation:** With the rapid uptake of WGS methods in surveillance and outbreak tracking, the findings obtained in this study can guide the selection of appropriate clustering methods for uncovering relevant transmission chains within a given time-period. For high resolution cluster analyses, WGS-SNP and cgMLST based analyses have similar clustering/timing characteristics even for data obtained from a high incidence setting.

## Research in context

### Evidence before this study

For nearly 30 years, molecular genotyping tools have been used to define transmission chains/clusters of *Mycobacterium tuberculosis* strains. A variety of tools are used for such analysis e.g. the presence/absence of spacers sequences (Spoligotyping), the length of tandem repeat patterns (24-loci-MIRU-VNTR) or, more recently, nearly the complete genome by whole genome sequencing (WGS). Each method has been proposed as the gold standard genotyping technique for detecting transmission events in a certain timeframe and selection of the optimal method for a given question is difficult as important parameters (e.g. the time span a particular outbreak can encompass) are not well defined. Based on inferred mutation rates, there have been some time scales proposed for clusters based on WGS SNP-based methods, supported by contact tracing data to confirm epidemiological links. However, there is uncertainty around these timing estimates for SNP-based techniques, limited timing estimates available for classical genotyping techniques and no such estimates for cgMLST approaches. This makes it very difficult for researchers, public health workers and clinicians to correctly interpret reported clustering data. This is especially the case as WGS based methods are becoming rapidly ingrained in surveillance and clinical workflows.

### Added value of this study

This study is the first to perform a comparative evaluation of cluster data defined by both classical and WGS-based *M. tuberculosis* genotyping approaches, especially with regard to transmission timing. While many studies have put forward various methods as the gold standard for *M. tuberculosis* transmission detection, we have tested clustering data generated by the different methods in a Bayesian statistical framework to elucidate the true fraction of recent transmission each approach is detecting. When specifically looking at recent transmission (e.g. less than 10 years previous), our results indicate that classical genotyping methods vastly over estimate recent transmission events. This solidifies the need for WGS-based methods when searching for recent outbreaks of *M. tuberculosis*.

### Implications of all the available evidence

Our study allows researchers and public health officials to select the appropriate genotyping method for assessing transmission with respect to the epidemiological setting and a given time-period. We also suggest the incorporation of particular genotyping methods in a cascade system with increasing resolution for various levels of surveillance e.g. from multi-country surveillance down to recent transmission and outbreak analyses. This is particularly important as each method comes with specific costs, infrastructure and computational requirements, human resources, and, last but not least interpretation complexities – all of which might not be feasible at any site or at any scale. Accordingly, our study can aid a cost/benefit analysis for selection of genotyping techniques in a cascade system, that might especially be used in high incidence, low resource settings.

## Introduction

Despite the large global efforts at curbing the spread of *Mycobacterium tuberculosis* complex (Mtbc) strains, 10.4 million new patients develop tuberculosis (TB) every year^1^. In addition, the prevalence of multidrug resistant (MDR) Mtbc strains is increasing^1^, predominantly through ongoing transmission within large populations^2,3^. The tracking and timing of recent transmission chains allows TB control programs to effectively pinpoint transmission hotspots and employ targeted intervention measures. This is especially important for the transmission of drug resistant strains as it appears that drug resistance may be transmitted more frequently than acquired^2^. Thus, interrupting transmission is key for the control of MDR-TB^3,4^. For the development of the most effective control strategies, there is a strong need for (i) appropriate identification of relevant transmission chains, risk factors and hotspots and (ii) robust timing of when outbreaks first arose.

Epidemiological TB studies often apply genotyping methods to Mtbc strains to determine whether two or more patients are linked within a transmission chain (molecular epidemiology)^5^. Contact tracing is the primary non-molecular epidemiological method for investigating transmission networks of TB, mainly based on patient interviews^6^. Although this method is often seen as a gold standard of transmission linking, it does not always match the true transmission patterns, even in low incidence settings^7^ and misses many connections^8^. The implementation of molecular genotyping and epidemiological approaches has overcome these limitations and is often used as the main approach for transmission analyses. Classical genotyping has involved *IS6110* DNA fingerprinting^9^, Spoligotyping^10^, and variable-number tandem repeats of mycobacterial interspersed repetitive units (MIRU-VNTR)^11^ which is the most common method at the moment^5^. The latter method is based on copy numbers of a sequence in tandem repeat patterns derived from 24 distinct loci within the genome^12^. If two patients have the same classical genotyping pattern such as a 24-loci MIRU-VNTR pattern (or up to one locus difference^12^) they are considered to be within a local transmission chain. The combination of spoligotyping and MIRU-VNTR-typing, where patterns must match in both methods to be considered a transmission link, is often considered the molecular gold standard for transmission linking and genotyping^12^. However, examples of unlinked patients with identical patterns have been observed, suggesting that this threshold covers too broad a genetic diversity and timespan between infections^7^.

The application of (whole genome) sequence (WGS)-based approaches for similarity analysis of Mtbc isolates and cluster determination is known to have high discriminatory power when assessing transmission dynamics^7,13–16^, either using core genome multi-locus sequence typing (cgMLST)^17,18^ or SNP distances^7,14,15,19^. WGS-based approaches compare the genetic relatedness of the genomes of the clinical strains under consideration, albeit usually excluding large repetitive portions of the genome, with the assumption that highly similar strains are linked by a recent transmission event^7,14^. Although many SNP cut-offs for linking isolates have been proposed^20^, the most commonly employed is based on the finding that a 5 SNP cut-off will cluster the genomes of strains from the majority of epidemiologically linked TB patients, with an upper bound of 12 SNPs between any two linked isolates^14^. The emerging widespread use of WGS has quickly pushed these cut-offs to be considered the new molecular gold standard of recent transmission linking, although SNP distances may vary for technical reasons (e.g. assembly pipelines or filter criteria^21^) and between study populations e.g. high and low incidence settings^19^.

In addition to cluster detection, uncovering the timing of transmission events within a given cluster is highly useful information for TB control e.g. for assessing the impact of interventions on the spread of an outbreak or uncovering when MDR-TB transmission first emerged in a particular setting. Accordingly, knowledge of the rate change associated with different genotyping methods is essential for correct timing. The whole genome mutation rate of Mtbc strains has been estimated by several studies as between 10^−7^ and 10^−8^ substitutions per site per year or ~0⋅3-0⋅5 SNPs per genome per year^7,14,22–24^, while the rate of change in the MIRU-VNTR loci specifically is known to be quicker (~10^−3^)^25^. Since these mutation rates have been shown to also vary by lineage^23,26^ and over short periods of time^22^, such variation needs to be accounted for when estimating transmission times, e.g. by using Bayesian phylogenetic dating techniques^3,22,25^.

Considering the multiple genotyping methods currently available, many of them proposed as a “gold standard”, there is an urgent need to precisely define the individual capacity of each method to accurately detect recent transmission events and perform timing of outbreaks. To provide this essential information, this study harnesses the power of WGS-based phylogenetic dating methods to assign timespans onto Mtbc transmission chains encompassed by the different genotypic clustering methods commonly used in TB transmission studies.

## Materials and Methods

### Dataset, ethical approval and sequencing

A set of 324 isolates from Kinshasa, Democratic Republic of Congo were collected from consecutive retreatment TB patients between 2005 and 2010 at TB clinics, servicing an estimated 30% of the population of Kinshasa. All isolates were phenotypically resistant to rifampicin (RR-TB) and the majority are also isoniazid resistant (i.e. MDR-TB). Use of the stored isolates without any linked personal information was approved by the health authorities of the DRC and the Institutional Review Board of the ITM in Antwerp (ref no 945/14). Libraries for whole genome sequencing were prepared from extracted genomic DNA with the Illumina Nextera XT kit, and run on the Illumina NextSeq platform in a 2×151bp run according to manufacturer’s instructions. Illumina read sets are available on the ENA (https://www.ebi.ac.uk/ena) under the accession number PRJEB27847.

### Genome reconstruction

The MTBseq pipeline^27^ was used to detect the SNPs for each isolate using the H37Rv reference genome (NCBI accession number NC000962.3)^28^. Sites known to be involved in drug resistance (as outlined in the PhyResSE list of drug mutations v27^29^) were excluded from the alignment and additional filtering of sites with ambiguous calls in >5% of isolates and those SNPs within a 12bp window of each other was also applied.

### Transmission cluster estimation methods

Five standard transmission clustering approaches were chosen for comparison and analysis (four additional methods are outlined in the supplementary methods). Clustering due to convergent genotypic patterns were removed as outlined in the supplementary methods. For each method, the total SNP distances were calculated to investigate the range of variability encompassed within each cluster. Maximum SNP distances were derived from pairwise comparisons of isolates within the SNP alignment using custom python scripts. A clustering rate was calculated for each method using the formula (n_c_-c)/n, where n_c_ is the total number of isolates clustered by a given method, c is the number of clusters, and n is the total number of isolates in the dataset (n=324).

### Spoligotyping and MIRU-VNTR

Spoligotype patterns were obtained from membranes following the previously published protocol^10^. Isolates were said to be clustered if all 43 spacers matched. Genotyping by MIRU-VNTR was undertaken as previously described^12^. 2μl of DNA was extracted from cultures and amplified using the 24 loci MIRU-VNTR typing kit (Genoscreen, Lille, France). Analysis of patterns was undertaken using the ABI 3500 automatic sequencer (Applied Biosystems, California, USA) and Genemapper software (Applied Biosystems). Isolates were said to be clustered if all 24 loci matched. Mixed MIRU-VNTR patterns were observed in 18 isolates although this mixing was not observed in the WGS data, likely due to subculturing for sequencing.

### SNP cut-off clustering

In this study, we employed the widely used 5 SNP (proposed by Walker et al. ^14^ as the likely boundary for linked transmission) and 12 SNP cut-offs (proposed maximum boundary) for cluster definition. Additionally, we employed a lower cut-off of 1 SNP to look for clusters of very highly related isolates. Pairwise SNP distances were calculated between all isolates. A loose cluster definition was used, where every isolate in a cluster at most the SNP cut-off from at least 1 other isolate in the cluster.

### Estimation of transmission times

To estimate the age and timespan of potential transmission clusters, SNP alignments were created from the convergence-free version of the three primary clustering types: Spoligotyping, MIRU-VNTR, and 12 SNP clusters.

A Bayesian approach to transmission time estimation was then undertaken. The SNP alignments were created as above for the three high-level clustering types. Each cluster method alignment was separately input to BEAST-2 v2.4.7^30^ to create a time tree for those isolates. These phylogenies were built using the following priors: GTR+GAMMA substitution model, a log-normal relaxed molecular clock model to account for variation in mutation rates^31^ and coalescent constant size demographic model^32^, both of which have been found to be suitable for lineage 4 isolates in a previous study^19^. The MCMC chain was run six times independently per alignment with a length of at least 400 million, sampled every 40,000^th^ step (MIRU & Spoligotyping: 600 million; 12 SNP: 500 million;). A log normal prior (mean 1.5×10^−7^; variance 1⋅0) was used for the clock model to reflect the previously estimated mutation rate of *M. tuberculosis* lineage 4^7,14,22–24^, while allowing for variation as previously suggested^22^. A 1/X non-informative prior was selected for the population size parameter of the demographic model. Isolation dates were used as informative heterochronous tip dates and the SNP alignment was augmented with a count of invariant sites for each of the four nucleotide bases to avoid ascertainment bias^33^. Tracer v1.6 was used to determine adequate mixing and convergence of chains (ESS >150) after a 25% burn-in. The chains were combined via LogCombiner v2.4.8^30^ to obtain a single chain for each clustering type with high (>1000) effective sample sizes. The tree samples were combined in the same manner and resampled at a lower frequency to create thinned samples of (minimum) 20,000 trees. The algorithm for estimating the timespan of transmission events encompassed by each method is outlined in Supplemental Figure 1. Briefly, for each cluster created by the given method, we defined the MRCA node as the internal node that connects all taxa in that cluster. The youngest node was then defined as the tip that is furthest from this MRCA within the clade (i.e. the tip descendant from that node that was sampled closest to the present time). To better account for changes in the mutation rate over short periods^22^, all trees estimated and sampled during the Bayesian MCMC process were used instead of only a single summary phylogeny. For each retained tree in the MCMC process, the difference in age between the MRCA node and youngest node was calculated. This gave a distribution of likely maximum transmission event times within that cluster. For each method, these per-cluster aggregated ages were then combined across all clusters to give a per-method distribution of transmission event times represented by the clusters. The 95% Highest Posterior Density (HPD) interval of these distributions was calculated with the LaplacesDemon p.interval function in R v3.4.0 and the distribution within this interval for each method along with the mean based upon this interval were then visualized in violin plots per clustering method using ggplot2 in R.

**Figure 1:**
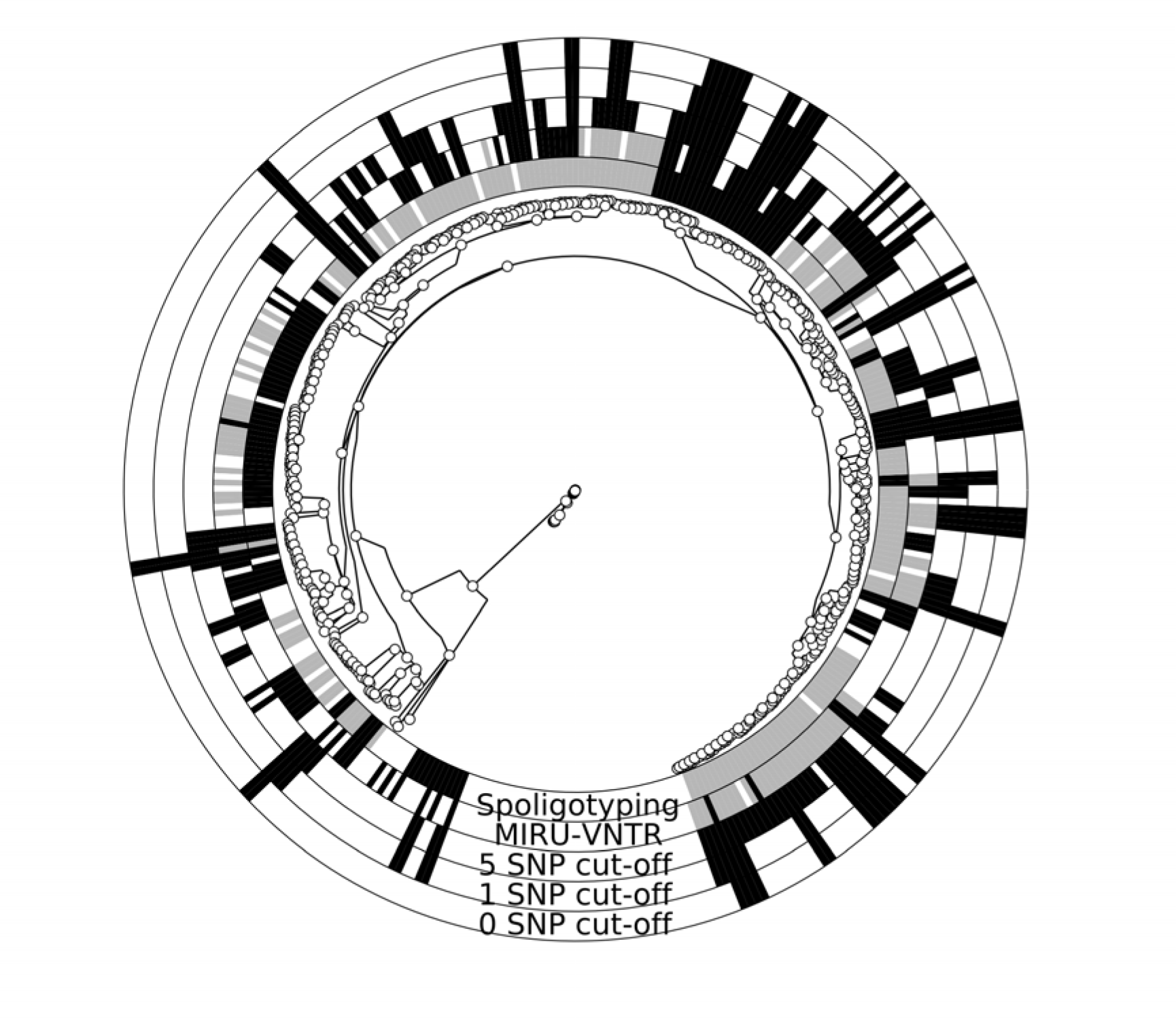
Clustering of *M. tuberculosis* isolates. For each approach the inclusion of an isolate into a cluster is outlined in the surrounding circles using GraPhlAn^45^. If an isolate is in a cluster not affected by convergence, it is highlighted in black for the given method. If an isolate is in a cluster affected by convergence, it is shown in grey. The ML phylogenetic tree is rooted between L4 and L5 isolates.

## Results

In this study, we assessed five different approaches for generating putative *M. tuberculosis* transmission clusters: Spoligotyping, MIRU-VNTR, and SNP-based clustering using a 12, 5 and 1 SNP cut-off, using a dataset of 324 isolates collected 2005-2010 in Kinshasa, Democratic Republic of Congo (DRC). The dataset contained 309 L4 and 15 L5 isolates, with a maximum of 1671 SNPs between any two isolates. Bayesian phylodynamic dating approaches implemented in BEAST-2^30^ were then utilised to assign timespans to the transmission events estimated by each genotyping method.

As expected, classical genotyping methods clustered many strains, with the lowest resolution (i.e. highest clustering rate) (Figure 1, Table 1). Convergent evolution (defined as the same pattern observed in unrelated strains; see supplemental methods) was found to affect 39% (12) of Spoligotyping-based clusters. Although MIRU-VNTR performed far better than spoligotyping, 16% (6) of clustering patterns were influenced by convergence in this study (Table 1, Figure 1). Spoligotyping and MIRU-VNTR patterns that were affected by convergence were removed from further timespan analyses. WGS-based methods had by far the highest discriminatory power and low SNP cut-offs grouped isolates into smaller clusters (e.g. 2-10 isolates per cluster for a 5 SNP cut-off) (Table 1, Figure 1). The high percentage of strains in a 12 SNP cluster (75%) suggests high levels of transmission in this population, making is suitable for further transmission analyses.

**Table 1:** Clustering method overview. For each clustering method, the general features are outlined in the table. Median ages and 95% HPD ranges are based upon the BEAST2 estimates of clade heights (see methods).

| Method | Strains in clusters | Number of clusters | Percent of strains in clusters | Cluster sizes | Maximum SNP distances | Clustering rate | Median Timespan | Timespan 95% HPD |
| --- | --- | --- | --- | --- | --- | --- | --- | --- |
| Spoligotype | 118 | 21 | 36.42 | 2-28 | 0-189 | 0.2994 | 76.51 | 0.81 - 823.21 |
| MIRU-VNTR | 121 | 32 | 37.35 | 2-11 | 0-48 | 0.2747 | 26.08 | 0 - 162.27 |
| 12 SNP cluster | 242 | 47 | 74.69 | 2-34 | 0-23 | 0.6019 | 23.63 | 0 - 102.58 |
| 5 SNP cluster | 147 | 40 | 45.37 | 2-27 | 0-10 | 0.3302 | 10.86 | 0 - 47.07 |
| 1 SNP cluster | 74 | 29 | 22.84 | 2-6 | 0-2 | 0.1389 | 3.91 | 0 - 23.54 |

Bayesian phylogenetic dating of the timeframe associated with particular transmission chains showed large differences in estimated cluster ages between the different genotyping approaches used (Table 1, Figure 2), correlating well with the difference in discriminatory power. Cluster ages are defined here as the most ancient transmission event that links any two isolates within a specific cluster (see methods and supplementary figure 1). Thus, in phylogenetic terms, the cluster age is the difference in time between when the most recent common ancestor (MCRA) of the entire cluster existed and the date of isolation of the furthest isolate from this ancestor.

**Figure 2:**
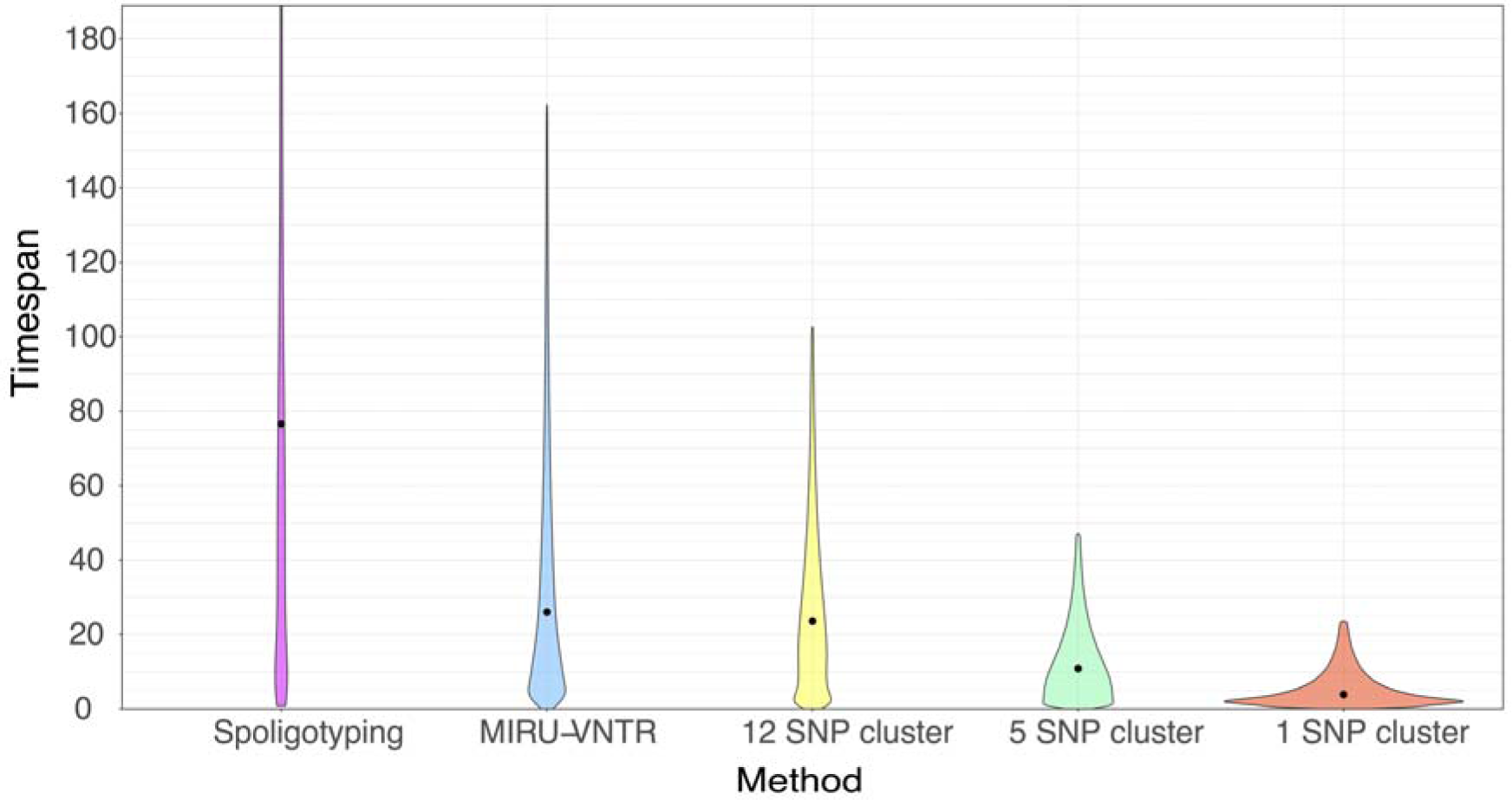
Timespans associated with transmission clusters. For each clustering method, the timespan associated with a cluster was estimated using BEAST-2. The ages of each cluster (Y-axis) was aggregated per clustering method (X-axis). Violin plots show the mean (black dot) for timespans along with the proportion of clusters with a given age (coloured kernel plots). Note the y-axis is cut at 200 years. See Table 1 for full 95% HPD for each method.

The aggregate median ages of clusters derived from Spoligotyping were found to often be several decades old (median 77 years (95% HPD: 1-823)) (Table 1, Figure 2). MIRU-VNTR clustering encompassed more recent transmission events than Spoligotyping, but were still found to be often over 2 decades old (median 26 years (95% HPD: 0-162)). The combination of MIRU-VNTR and Spoligotyping resulted in cluster ages similar to MIRU-VNTR alone (supplemental table 1). Clusters based on SNP cut-offs correlated to 23 years using a 12 SNP cut-off (95% HPD: 0-103), 11 years using a 5 SNP cut-off (95% HPD: 0-47), and 4 years using a 1 SNP cut-off (95% HPD: 0-24) (Table 1, Figure 2). For comparison, the cgMLST method was also performed and discriminatory power, cluster sizes and ages based on cgMLST alleles were similar to the SNP-based clusters (see supplemental methods, supplemental table 1).

## Discussion

The term ‘recent transmission’ is often applied to gain a better understanding of the current transmission dynamics of pathogens in a given population. However, little data is available on how recent a likely transmission event occurred when measured with different genotyping methods. To get a better understanding of the discriminatory power of different classical genotyping techniques and WGS-based approaches in relation to outbreak timing, this study has performed an in-depth comparison of clustering rates and dated phylogenies obtained in a collection of 324 Mtbc strains from a high incidence setting (Kinshasa, DRC). With a whole genome phylodynamic approach employed as a gold standard, our study demonstrates that each genotyping method was associated with a specific discriminatory power resulting in clusters representing vastly different time periods of transmission events (Table 1 and Figure 2). This has significant implications for data interpretations e.g. when selecting and utilising different genotyping/clustering approaches for epidemiological studies and assessing the effectiveness of public health intervention strategies.

As the most extreme example, Spoligotyping-derived clusters were associated with transmission events that can be several decades old. This low discriminatory power coupled with the high rate of convergent evolution (the same spoligotype pattern found in phylogenetically distant isolates) adds weight to the previous suggestion that this technique is not suitable for recent transmission studies^34^, although may be of use as a low-cost method of sorting Mtbc strains into the seven primary lineages^35,36^.

In line with previous findings^34,37^, convergent evolution of 24-loci MIRU-VNTR patterns was rarer than observed for Spoligotyping, but did occur in 16% of MIRU-VNTR-based clusters. Additionally, the transmission times encompassed by MIRU-VNTR clusters often spanned over two decades (Table 1, Figure 2), confirming previous studies showing over-estimation of recent transmission with this method^7,13,19,38^.

For defining transmission events that occurred in more recent time frames before sampling, WGS-based methods were found to be better suited than classical genotyping methods (Table 1, Figure 2). The 12 SNP cut-off, currently the recommended upper bound for clustering isolates, often defines transmission events that occurred on average two decades prior to sampling, similar in age to clusters estimated by MIRU-VNTR. This suggests that the 12 SNP cluster method may be a good replacement for MIRU-VNTR as it detects larger transmission networks spanning similar transmission time periods but is less affected by convergent evolution. Isolates clustered at a low (5 SNP) or nearly identical (1 SNP) cut-off were found to represent transmission events occurring over a time span of up to ten years. These findings correlate well with previous studies where confirmed contact tracing-based epidemiological links were found between patients that were two^15,39^ and three^7^ SNPs apart. The original paper that proposed the 5 and 12 SNP cut-offs found that serial isolates that were 10 years apart differed by, on average, 6 SNPs, also agreeing with the findings presented here ^14^. Comparisons between the SNP-based (using almost all genomic differences) and the cgMLST-based (using a defined core set of genes) methods demonstrated that the latter approach gives similar estimates to full SNP approaches. This supports the use of low SNP or cgMLST differences for detection or exclusion of very recent transmission, although this low variability between isolates makes robust identification of transmission direction difficult, especially during short timespans.

The mutation rate of *M. tuberculosis* has been estimated to be between 10^−7^ and 10^−8^ substitutions per site per year^3,7,23^. Within the Bayesian analysis employed here, the mutation rate was free to vary between these values but was found to strongly favour ~3×10^−8^ (ESS > 1000 for all runs; 95% HDP: 4×10^−9^ – 8×10^−8^), translating to approximately 0⋅3 SNPs per genome per year. While the mutation rate used here is in line with previous estimates for lineage 4 ^23^ (which most of this dataset is comprised of), it may be similar in other lineages, although this has only been shown for lineage 2^3,23^. Thus, per-lineage estimates are required for all seven lineages to ensure similar transmission times are linked to genotyping methods across the whole diversity of the Mtbc.

While this study has many advantages due to its five year population based design in an endemic setting coupled with the application of three different genotyping methods (Spoligotyping, 24-locus MIRU-VNTR and WGS), future confirmatory studies could address the following drawbacks that are inherent to genomic epidemiology^16,21^: 1) studies employing contact tracing and/or digital epidemiology^40^ in conjunction with these genotyping methods can help confirm transmission times associated with different clusters (although these methods also have many limitations); 2) as outlined above, strains of other lineages of the Mtbc should be analysed in a similar fashion to ensure transferability of findings across the entire complex; 3) a broad range of drug resistance profiles should be included to fully assess the impact of such mutations on transmission estimates; 4) improved WGS methods, such as directly from clinical samples to help reduce culture biases^41^ and longer reads (e.g. PacBio SMRT or Nanopore MinION) to capture the entire genome, including repetitive regions such as PE/PPE genes known to impact genome remodelling^42,43^, will ensure that the maximum diversity between isolates is captured; 5) extensive panels of Spoligotyping and MIRU-VNTR results paired with WGS data will help assess the extent of convergence in these methods and better correlate their clusters with those of low SNP thresholds and 6) standardised SNP calling pipelines appropriate across all lineages, with high true positive/low false negative rates, will ensure that Mtbc molecular epidemiology can be uniformly implemented and comparable across studies.

Since each method was found to represent different timespans and clustering definitions, they can be used in a stratified manner in an integrated epidemiological and public health investigation addressing the transmission of Mtbc strains. For instance, although Spoligotyping clusters represented potentially very old transmission events, the low associated cost and its ability to be applied directly on sputum helps reduce culture bias and thus robustly assign lineages. Thus, Spoligotyping and/or MIRU-VNTR would serve well as first-line surveillance of potential transmission events in the population, guiding further investigations and resource allocations.

These potential transmission hotspots could be further investigated with contact tracing and/or WGS. Employment of different cut-offs and clustering approaches to WGS data can then address several questions. The 12 SNP/cgMLST allele cluster approaches serve well for high level surveillance targeting larger (older) transmission networks, akin to what is currently often done using MIRU-VNTR (e.g.^15,44^). Recent transmission events can then be detected through employment of low SNP cut-offs (e.g. 5 SNPs for transmission in the past 10 years or 1 SNPs for transmission in the past 5 years). In high incidence/low diversity settings where amalgamation of clusters may inadvertently obscure distinct hotspots of transmission at different time points, subdivision into distinct time-dependant clusters can be undertaken using the algorithm presented in such a study in East Greenland^19^.

Overall, phylodynamic approaches applied to whole genome sequences, as undertaken here, are recommended to fully investigate the specific transmission dynamics within a study population to account for setting-specific conditions, such as low/high TB incidence, low/high pathogen population diversity, sampling fractions and social factors influencing transmission. As WGS methods become more commonplace and easier to implement in a variety of settings, each genotyping method can be employed as part of an overall evidence gathering program for transmission, placing molecular epidemiological approaches as an integral part in tracking and stopping the spread of TB.

## Acknowledgements

The authors would like to thank Armand Van Deun and Koen Vandelannoote for valuable discussion and input and Cecile Uwizeye for aid with spoligotyping.

## Author contributions

CJM, FG and BCdJ conceived the study. MKK and BCdJ oversaw collection of isolates and ethical approval. TAK, SA, MM, PB and SN undertook classic genotyping and sequencing of isolates. CJM, PM, TA, CU and PL undertook WGS assembly and data preparation. CJM undertook all convergence and clustering analyses. CJM, PM, JP, MM, TS and DK undertook all phylodynamics. CJM, PM, SN and BCdJ wrote the manuscript. All authors read and revised the manuscript and approved its final form.

